## Supplementary for "Climate-induced range shifts drive adaptive response via spatio-temporal sorting of alleles"

#### CONTENTS

**S1 Supplementary Methods** **2**

S1.1 **Study populations and sequencing protocol** 2

S1.2 **Population genetic structure and biogeographic barriers** 3

S1.3 **Ancestral sequence reconstruction** 4

S1.4 **Expansion signal** 5

S1.5 **Demographic inference** 6

S1.6 **Distribution modelling** 7

S1.7 **Visualising shifts in environment space and habitat availability** 7

S1.8 **Predicting adaptive genetic structure in space and time** 7

S1.9 **Validation of glacial genomic offset** 8

S1.10 **References** 9

**S2 Supplementary Figures** **11**

**S3 Supplementary Tables** **28**

### S1 Supplementary Methods

#### S1.1 Study populations and sequencing protocol

170 natural populations of *Dianthus sylvestris* were sampled between 2008-2018 for leaf material, which were dried and stored in silica gel. Of these, we selected 116 populations that covered the geographic range and ecological diversity of the species, and 5-20 individuals per population, for DNA extraction and sequencing (sample map in **Fig. 1**). DNA extraction and library preparation followed a modified protocol[1] of the Illumina Nextera DNA library preparation kit. Individuals were pooled into libraries and indexed with unique dual-indexes from Integrated DNA Technologies Co, to avoid index-hopping[2]. Four libraries totalling 1261 individual samples were sequenced (150bp paired-end sequencing) in four lanes of an Illumina NovaSeq 6000 machine at Novogene Co, resulting in a coverage of ca. 2x per individual. We processed demultiplexed raw sequencing reads as detailed below, taking steps to i) correct for biases associated with low-coverage data (e.g. recalibration of base quality scores) and ii) minimise information loss given that we account for statistical uncertainties by working directly on genotype likelihoods in downstream analyses.

Raw reads were trimmed with *Trimmomatic* version 0.35[3] to remove library (Illumina Nextera) adapters; allowing for a maximum mismatch of 2 bases, a palindrome clip threshold of 30 bases, a simple clip threshold of 10 bases, a minimum adapter length of 2 bases, and keeping both reads. Trimmed paired-end reads were mapped against the reference genome (Fior et al., paper in preparation) via the BWA-MEM algorithm of *BWA* version 0.7.17[4,5] using default parameters. Mapped SAM files were converted to binary format (BAM) and sorted with *Sambamba* version 0.6.8[6]. Reads with mapping quality of < 20 were filtered out. PCR duplicates were removed using MarkDuplicate in *Picard Toolkit* version 2.0.1 (http://broadinstitute.github.io/picard). Local-realignment of indels was performed across all 1261 samples using *GATK* version 3.5[7], via the tools RealignerTargetCreator and IndelRealigner. The former generated indel targets prior to realigning indels using the latter. Samples' base quality scores were recalibrated using *ATLAS* version 0.9[8], to correct for biases that affect low-coverage sequencing data[9]. Specifically, we estimated recalibration parameters for each set of samples belonging to a sequencing lane-run combination (4 sets of recalibration parameters in total); restricting the analyses to sites covered at least twice, assuming equal base frequencies, and employing the 'qualFuncPosSpecificContext' model. Recalibration required a set of sites known or inferred to be monomorphic. We acquired such a set from an independent sequencing run of 192 *D.sylvestris* individuals from 2017, where we inferred positions that are highly conserved among populations by removing all potentially polymorphic sites. After estimation of lane-specific recalibration parameters, individual BAM files were recalibrated. Finally, we clipped overlapping read pairs using *bamUtil* version 1.0.14[10] clipOverlap.

#### S1.2 POPULATION GENETIC STRUCTURE AND BIOGEOGRAPHIC BARRIERS

To perform principle component analysis (PCA) and admixture analysis on low-coverage samples, we used *PCAngsd* version 0.98[11]. PCA was performed on both the full dataset (of 1261 individuals) and on a balanced dataset comprising a common, down-sampled size of 125 individuals per geographic region, while admixture was analysed exclusively on the balanced dataset. The use of the balanced dataset was to account for known biases related to uneven sampling across clusters[12–15], and was generated by selecting 125 individuals from 22-25 populations (5-10 individuals per population) that evenly covered each of the three geographic regions (**Supplementary Table S1**). To run *PCAngsd*, we first generated genotype likelihoods in *ANGSD* version 0.933[16] for a set of samples and outputted them in Beagle format. We employed the GATK (-GL 2) genotype likelihood model in *ANGSD*. We filtered out flagged sites (i.e. not primary, failure and duplicate reads) and sites with mapping or base quality scores below 1. We reduced the effect of reads with excessive mismatches (-C 50) and only retained pairs of reads with both mates mapped correctly (i.e. proper pairs). We performed PCA on the resultant Beagle genotype likelihood file using *PCAngsd* under the default settings. We estimated individual admixture proportions under *PCAngsd*, using the admix argument (-admix) and iterating over K's (K = 2-6) via the -e argument which defines the number of eigenvalues. We automatically searched for the optimal alpha (i.e. the sparseness regularisation parameter) specifying a soft parameter upper-bound of 10,000.

To test if admixture (as inferred by *PCAngsd*) reflects true (biological) admixture or is an artefact of demographic scenarios such as recent bottlenecks that may violate method assumptions[17], we used chromosome painting and patterns of allele sharing to construct painting palettes via the programs *MixPainter* and *badMIXTURE* (using the unlinked model)[17]. These painting palettes were then compared to the *PCAngsd*-inferred admixture palettes. We first imputed and phased the data using *Beagle* version 3.3.2[18]. Different Beagle inputs files comprising different sets of individuals were generated depending on the pair of lineages to assess (see Methods and Table S1 for details). We converted the Beagle output to Chromopainter input format using an accessory script included with the program and constructed painting palettes with MixPainter[17].

To construct a genetic distance tree, we calculated pairwise genetic distances between 549 individuals (5 individuals per population for all populations; **Supplementary Table S1**) using *ATLAS* version 1.0[8]. We first generated individual genotype likelihood files (GLFs) from BAM files. We then calculated genetic distances between all individual pairwise combinations, employing a distance measure (or weight) reflective of the number of alleles differing between the genotypes (distType=squaredDiff).

To estimate population pairwise $F_{ST}$, we relied on 5 individuals per population, to allow standardised comparison across all population-pairs. We first estimated site allele frequency likelihoods based on individual genotype likelihoods in *ANGSD* version 0.933[16]. We used the GATK (-GL 2) genotype likelihood model in *ANGSD*, as we noticed irregularities in the SFS with the SAMtools (-GL 1) model similar to that reported in Korneliussen et al.[16] with particular BAM files. We discarded sites with global sequencing depths of below 10 and above 75; i.e. to necessitate an average minimum coverage of 1x per haploid chromosome and to remove sites that are excessively represented which may be indicative of paralogs. We filtered out flagged sites (i.e.

not primary, failure and duplicate reads) and sites with mapping or base quality scores below 1. We retained sites where the individual sequencing depth was at least 1x from at least 3 individuals, kept only pairs of reads with both mates mapped correctly (i.e. proper pairs), and adjusted mapping qualities for reads with excessive mismatches (-C 50). Using these population site allele frequency likelihoods, we generated maximum likelihood estimates of (folded) 2-population site frequency spectra (2D-SFS) via the *realSFS* function in *ANGSD*. Using the 2D-SFS as a prior in conjunction with each populations' site allele frequency likelihoods, we then estimated $F_{ST}$ using Bhatia's estimator[19], which is preferable for small samples sizes.

#### S1.3 Ancestral sequence reconstruction

The phylogenetic tree used in ancestral sequence reconstruction was taken from Fior et al. (paper in preparation), who inferred a comprehensive maximum likelihood phylogenetic tree of 30 European taxa of *Dianthus*. The subset topology relevant and used in ancestral sequence reconstruction is described in Newick format below:

((((((*D.lusitanus*,*D.pungens*),(*D.superbus_alpestris*,*D.superbus_superbus*)),(*D.sylvestris_alpine*,*D.sylvestris_apennine*)),*D.carthusianorum*),*D.glacialis*),*D.deltoides*)

For ancestral sequence reconstruction, we produced paired-end libraries for one individual per species representing the phylogenetic tree indicated above using the Illumina TruSeq DNA library preparation kit. Libraries were sequenced on an Illumina HiSeq 4000 machine at the Functional Genomics Center Zurich (FGCZ, Switzerland). This generated a coverage of ca. 9-10x per individual. Raw sequencing reads were trimmed with *Trimmomatic* version 0.35[3] to remove library (Illumina TruSeq) adapters; allowing for a maximum mismatch of 2 bases, a palindrome clip threshold of 30 bases, a simple clip threshold of 10 bases, a minimum adapter length of 2 bases, and keeping both reads. Low quality bases (≤ 3) from the leading and trailing ends of the read were trimmed, reads were clipped when the average quality within a 4bp sliding window fell below 15, and reads were removed if shorter than 36 bp. Trimmed paired-end reads were mapped against the *D. sylvestris* reference genome via the BWA-MEM algorithm of *BWA* version 0.7.17[4,5] under default settings, filtered for a mapping quality of < 20, and had PCR duplicates removed via *Picard Toolkit* version 2.0.1 (http://broadinstitute.github.io/picard). Overlapping read pairs were clipped using *bamUtil* version 1.0.14[10] clipOverlap.

To generate FASTA sequences for the different species, we used *GATK* version 3.5[7] FastaAlternateReferenceMaker. This tool replaces reference bases at variation sites with bases defined in a given variant callset, while retaining reference bases elsewhere. Additionally, a mask can be applied such that select sites, e.g. ambiguous calls, can be set to 'N' in the resultant FASTA. We generated species-specific variant callsets with *freebayes* version 1.3.1[20], reporting records for every position in the genome including monomorphic sites and sites with zero depth, as well as information for all haplotype alleles. We used default parameters to call variants, i.e., variants needed to be supported by ≥ 2 observations in a single sample and by ≥ 20% of the reads from a single sample. We partitioned the resultant genomic VCF into four partitions (using *bcftools* version 1.8[21]):

- Partition I: Sites that are confidently inferred as variants. Here, we first extracted potential variant sites from the genomic VCF (i.e. sites with > 1 alternate allele). We retained variants with a minimum quality score of 20 and a minimum depth of 3.
- Partition II: Monomorphic sites that are inferred as conserved i.e., identical to the *D. sylvestris* reference.
- Partition III: Sites whose variant status is unresolved. These are variant sites that did not pass the filtering in Partition I.
- Partition IV: Sites with zero depth.

We then used FastaAlternateReferenceMaker to replace the reference bases at variation sites with species-specific variants (Partition I), while masking (i.e. setting as "N") sites under Partitions III and IV. Species FASTA files were subsequently combined into a multi-sample FASTA. Using this, we probabilistically reconstructed ancestral sequences at ancestral nodes via *PHAST* version 1.4 prequel[22], using a tree model produced by *PHAST* phylofit under a REV substitution model and a tree topology specified by prior phylogenetic work on this clade (Fior et al., paper in preparation). Ancestral sequence FASTA files were then generated for each ancestral node (of the phylogenetic tree) from the prequel results using a custom script (PHAST2fasta.py, available at: https://github.com/hirzi/RhEA).

#### S1.4 Expansion signal

We calculated the population pairwise directionality index $\psi$ using equation 1b from Peter and Slatkin[23], shown below:

$$\psi = \sum_{i=1}^{n} \sum_{j=1}^{n} (i - j) f_{ij}$$

where $f_{ij}$ is the fraction of SNPs in the sample that are at frequency $i$ in population 1 and at frequency $j$ in population 2, with the SFS normalized such that:

$$\sum_{i=1}^{n} \sum_{j=1}^{n} f_{ij} = 1$$

Two-population site frequency spectra (10 individuals per population) for calculating ψ were estimated in *ANGSD* and *realSFS*[24]. Unfolding of spectra was achieved via polarisation with respect to the ancestral state of sites defined at the *D. sylvestris* (Apennine lineage) - *D. sylvestris* (Alpine lineage) ancestral node. To estimate 2D-SFS, we first estimated populations' site allele frequency likelihoods in *ANGSD*, under the GATK (-GL 2) genotype likelihood model. We discarded sites with global sequencing depths of below 20 and above 150; i.e. to necessitate an average minimum coverage of 1x per haploid chromosome and to remove sites that are excessively represented which may be indicative of paralogs. We filtered out flagged sites (i.e. not primary, failure and duplicate reads) and sites with mapping or base quality scores below 1. We retained sites where the individual sequencing depth was at least 1x from at least 3 individuals, kept only pairs of reads with both mates mapped correctly (i.e. proper pairs), and adjusted mapping qualities for reads with excessive mismatches (-C 50). Using these population site allele frequency likelihoods, we generated maximum likelihood estimates of the 2D-SFS via *realSFS*.

#### S1.5 DEMOGRAPHIC INFERENCE

For our distance-based species phylogeny, we combined the TruSeq sequencing dataset (9 species-specific samples; **Supplementary Methods 1.3**) with low-coverage (ca. 2x) whole genome data from 1 representative each of the three *D. sylvestris* lineages (Alpine, Apennine and Balkan). To avoid biases related to unequal coverage across samples in the calculation of genetic distances, we down-sampled all samples to ca. 2x via *SAMtools*[21]. Note that our limited representation of species across the *Dianthus* clade and reliance on low-coverage data here motivated the use of the topology from Fior et al. (paper in preparation) for the ancestral sequence reconstructions, while our purpose here is to acquire a rooted topology for the *Dianthus sylvestris* sub-clade. To estimate distance-based phylogenies on this combined dataset, we used *ngsDist*[25] that accommodates genotype likelihoods in the estimation of genetic distances. We first produced the input file for *ngsDist*, i.e. genotype likelihoods in Beagle format, using *ANGSD* under the GATK (-GL 2) genotype likelihood model. We filtered out flagged sites (i.e. not primary, failure and duplicate reads) and sites with mapping or base quality scores below 20. We retained sites where the individual sequencing depth was at least 1x from at least 3 individuals, kept only pairs of reads with both mates mapped correctly (i.e. proper pairs), and adjusted mapping qualities for reads with excessive mismatches (-C 50). Genetic distances were then calculated in ngsDist[25] i) genome-wide and ii) along 10kb windows (see Methods).

For demographic inference, we estimated unfolded three-population site frequency spectra (3D-SFS) comprising one representative population each per lineage via *ANGSD* and *realSFS*. The three representative populations were Kapetovo Jezero (Balkan), Civita (Apennine) and Polsa (Alpine) (**Supplementary Table S1**). Based on LGM hindcasts of lineage distribution models, these populations are inferred to have been relatively stable over the last 20kya, and, importantly, not products of recent post-glacial expansion. To calculate 3D-SFS, we used 20 individuals per population and first estimated populations' site allele frequency likelihoods in *ANGSD*, under the GATK (-GL 2) genotype likelihood model. We discarded sites with global sequencing depths of below 40 and above 300; i.e. to necessitate an average minimum coverage of 1x per haploid chromosome and to remove sites that were excessively represented and potentially indicative of paralogs. We filtered out flagged sites (i.e. not primary, failure and duplicate reads) and sites with mapping or base quality scores below 1. We retained sites where the individual sequencing depth was at least 1x from at least 5 individuals, kept only pairs of reads with both mates mapped correctly (i.e. proper pairs), and adjusted mapping qualities for reads with excessive mismatches (-C 50). Using these population site allele frequency likelihoods, we generated maximum likelihood estimates of the 3D-SFS via *realSFS* (100 bootstrap replicates), polarising the spectra with respect to the ancestral state of sites defined at the *D. lusitanus* - *D. sylvestris* (Alpine lineage) ancestral node (for tree topology, see **Supplementary Methods S1.3**).

Demographic parameter estimation was then performed in *moments*[26], for a set of 18 models (**Supplementary Fig. S10**). To facilitate model selection and optimisation in moments, we employed an iterative optimisation procedure (modified from Portik et al.[27]) that started with the generation of 10 sets of threefold randomly perturbed parameters. Each parameter set was optimised using the Nelder–Mead method (optimize_log_fmin), and ran for maximum of 3 iterations per algorithm step. The 3D-SFS was simulated for each optimised parameter set, and the likelihood of the data given the simulated spectrum was estimated using a multinomial approach[26] and recorded. The parameter set that generated the best-fit model (i.e. highest log-likelihood) was then used as starting values for a second round of optimisation, with twofold perturbed parameters, 20 replicates and 5 iterations per algorithm step. This iterative procedure was run for a further two rounds (four rounds total); with twofold parameter perturbation, 30 replicates and 10 iterations per algorithm step in the third round, and onefold parameter perturbation, 40 replicates and 15 iterations per algorithm step in the final round; to estimate demographic model parameters and final model log-likelihoods. Code used for running *moments* is available at: https://github.com/hirzi/RhEA.

#### S1.6 DISTRIBUTION MODELLING

Our R code for performing species distribution models is available at: https://github.com/hirzi/RhEA.

#### S1.7 VISUALISING SHIFTS IN ENVIRONMENT SPACE AND HABITAT AVAILABILITY

Our R code for visualising shifts in environment space and habitat availability through time is available at: https://github.com/hirzi/RhEA.

#### S1.8 PREDICTING ADAPTIVE GENETIC STRUCTURE IN SPACE AND TIME

To predict adaptive genetic variation in space and time, we modelled the association of genetic variants (SNPs) with changes in environment, using gradient forest (GF)[28,29]. For the response variable, our GF model takes population allele frequencies. Genetic variants (SNPs) segregating across 43 Alpine populations comprising 14 individuals each (602 individuals in total) were identified using the program *freebayes* version 1.3.1[20]. For variant-calling, we required sites to have a minimum global coverage of 602x (i.e. average 1x per diploid individual), limited (i.e. down-sampled) per-sample coverage to 15x, and skipped processing of alignments that overlapped positions with a global coverage ≥ 9030x (602 x 15). We excluded alignments from analysis if they had a mapping quality less than 1, and excluded alleles if their supporting base quality was less than 1. We required variants to be supported by at least 2 observations in a single sample and by ≥ 20% of the reads from a single sample. We calculated the marginal probability of genotypes, and reported genotypes using the maximum-likelihood estimate provided from genotype likelihoods.

Following variant-calling, we retained variants which lied exclusively in exon regions. We subsequently applied the following filtering strategy on identified exon variants, to retain a set of confidently-called variants: i) a minimum quality score of 30, ii) additional contribution of each observation ≥ 10 log units (Q10 per read), iii) variants observed on both (forward and reverse) strands, iv) variants observed in at least two reads "balanced" on each side of the site, v) sites represented in at least 90% of samples, vi) a minimum global read depth of 1204x (i.e. an average ≥ 1x per haploid sample), and vii) a minimum allele count of 14. We decomposed complex variant calls into phased SNP and INDEL genotypes, retaining, ultimately, only SNPs. All filtering was done using *bcftools* version 1.8.

Population allele frequencies were estimated from the resultant VCF genotype likelihoods using *vcflib* popStats version 1.0.1.1[30]. We filtered the population allele frequency dataset to retain only sites with depth ≥7 per population. GF was run on the resultant set of 390,262 SNP allele frequencies in batches of 10,000 SNPs, using 500 decision trees, and as outlined in Methods. Batch runs were combined via combinedGradientForest {standardize="after", method=2} in the *gradientForest* R package. Our code for running GF is available at: https://github.com/hirzi/RhEA.

#### S1.9 VALIDATION OF GLACIAL GENOMIC OFFSET

Population genetic diversity and neutrality statistics used for validating glacial genomic offsets were estimated in *ANGSD* version 0.933[16]. To estimate population genetic statistics, we first calculated site allele frequency likelihoods for each population (same populations and set individuals as in GF analysis above). We discarded sites with global sequencing depths of below 28 and above 210; i.e. to necessitate an average minimum coverage of 1x per haploid chromosome and to remove sites that are excessively represented which are potentially indicative of paralogs. We filtered out flagged sites (i.e. not primary, failure and duplicate reads) and sites with mapping or base quality scores below 1. We retained sites where the individual sequencing depth was at least 1x from at least 5 individuals, kept only pairs of read with both mates mapped correctly (i.e. proper pairs), and adjusted mapping qualities for reads with excessive mismatches (-C 50). Using these population site allele frequency likelihoods, we generated maximum likelihood estimates of the 2D-SFS using *realSFS* and from this, calculated population diversity and neutrality statistics, namely: nucleotide diversity $\pi$, Tajima's $D$, Fu & Li's $F$, Fay & Wu's $H$, and Zeng's $E$.

### S2 Supplementary Figures

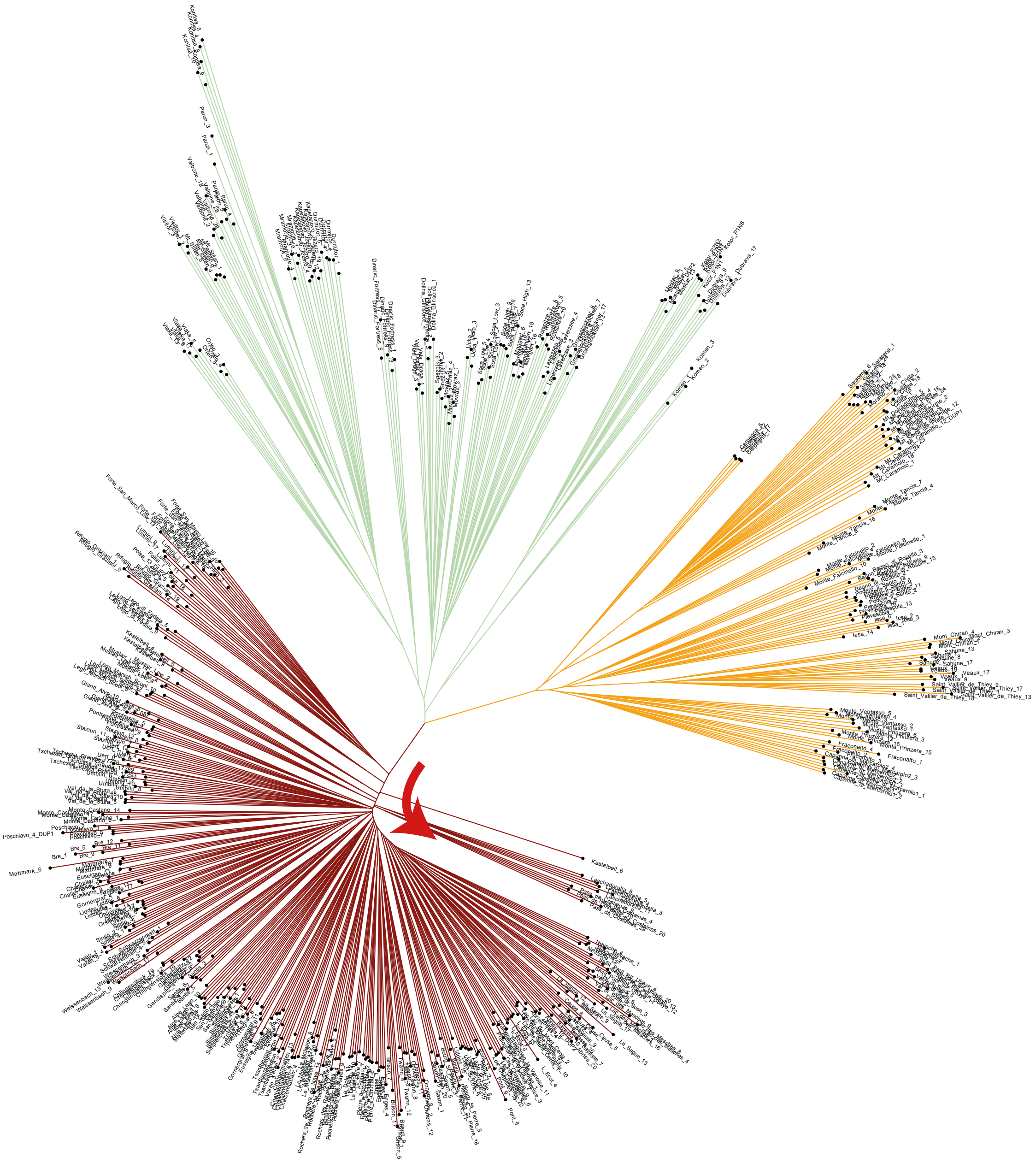

Figure S1. Distance-based phylogeny based on pairwise whole-genome genetic distances between 549 individuals (5 individuals per population for all sampled populations). Three distinct clades are evident, and are coloured by region-lineage (Alpine – dark red; Apennines –yellow; Balkans – green). The order of populations from central node to terminus within the Alpine clade (red arrow) in this tree closely follows an east to west geographic ordering, and alludes towards the direction of the post-glacial expansion of the Alpine lineage.

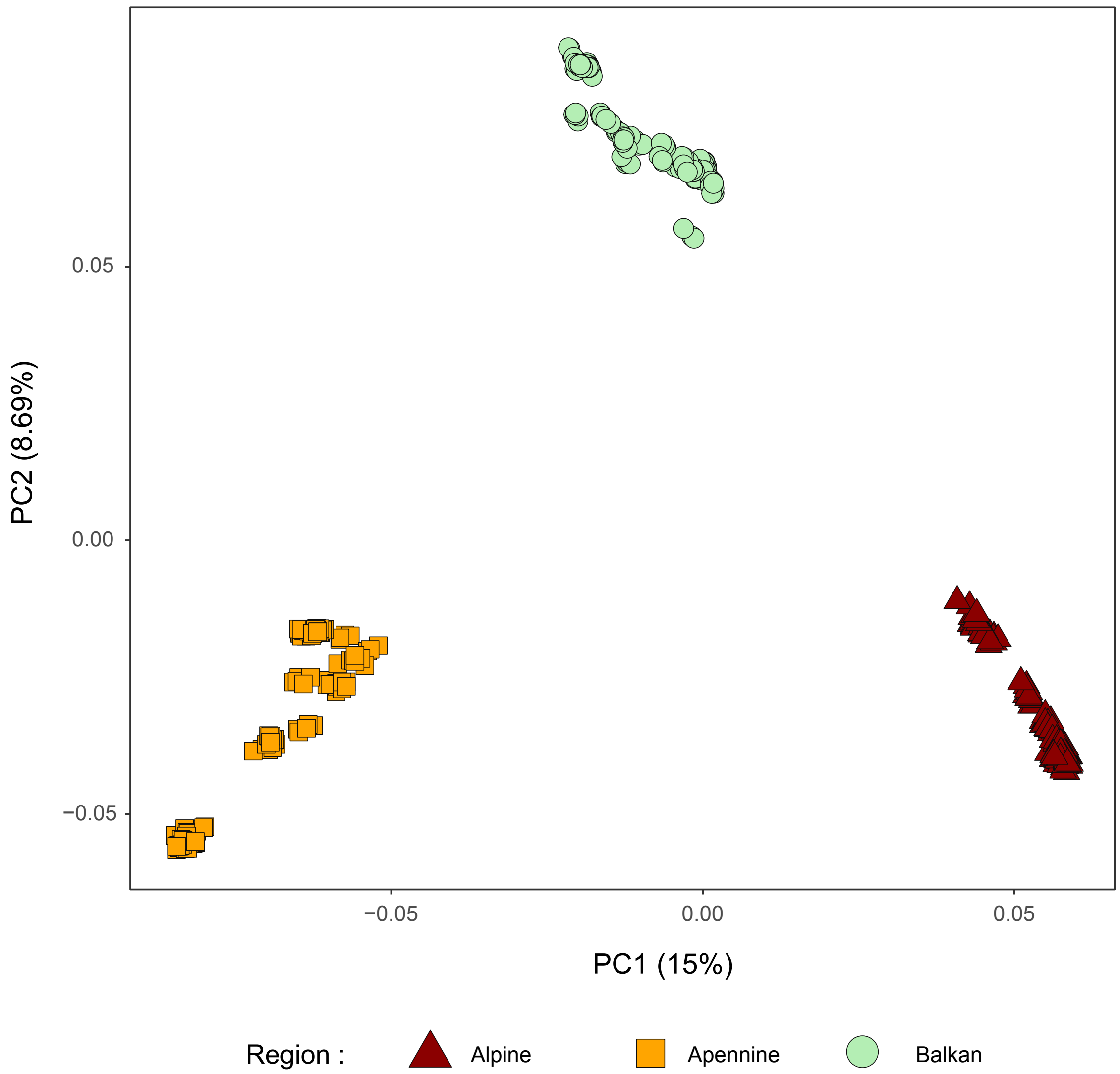


FIGURE S2. Principal component analysis (PCA) of whole-genome sequences, with eigenvalues in axes' labels. To account for effects of unbalanced sampling, the Alpine, Apennine and Balkan regions were down-sampled to 125 individuals each.

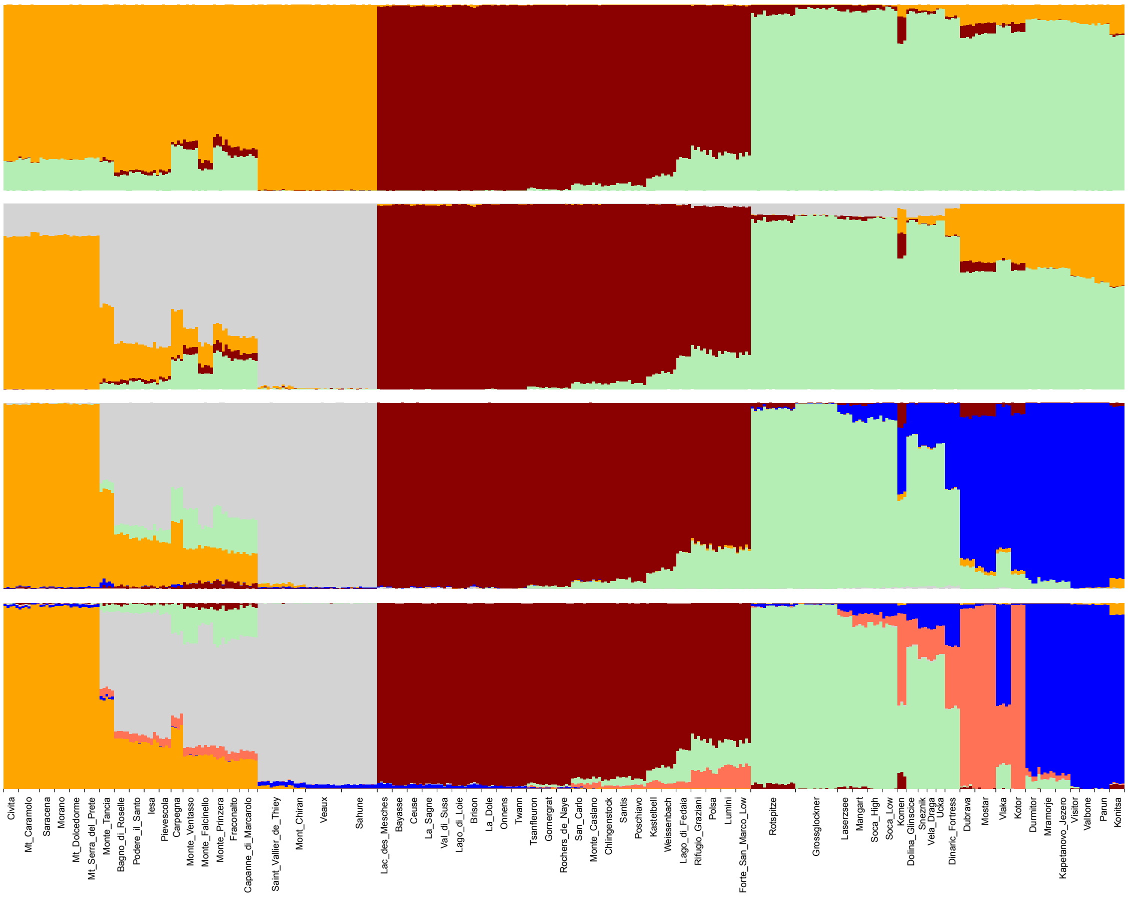


FIGURE S3. Admixture proportions of whole-genome sequences at K=3-6 (top to bottom). To minimise effects of unbalanced sampling, the Alpine, Apennine and Balkan regions were down-sampled to 125 individuals each.

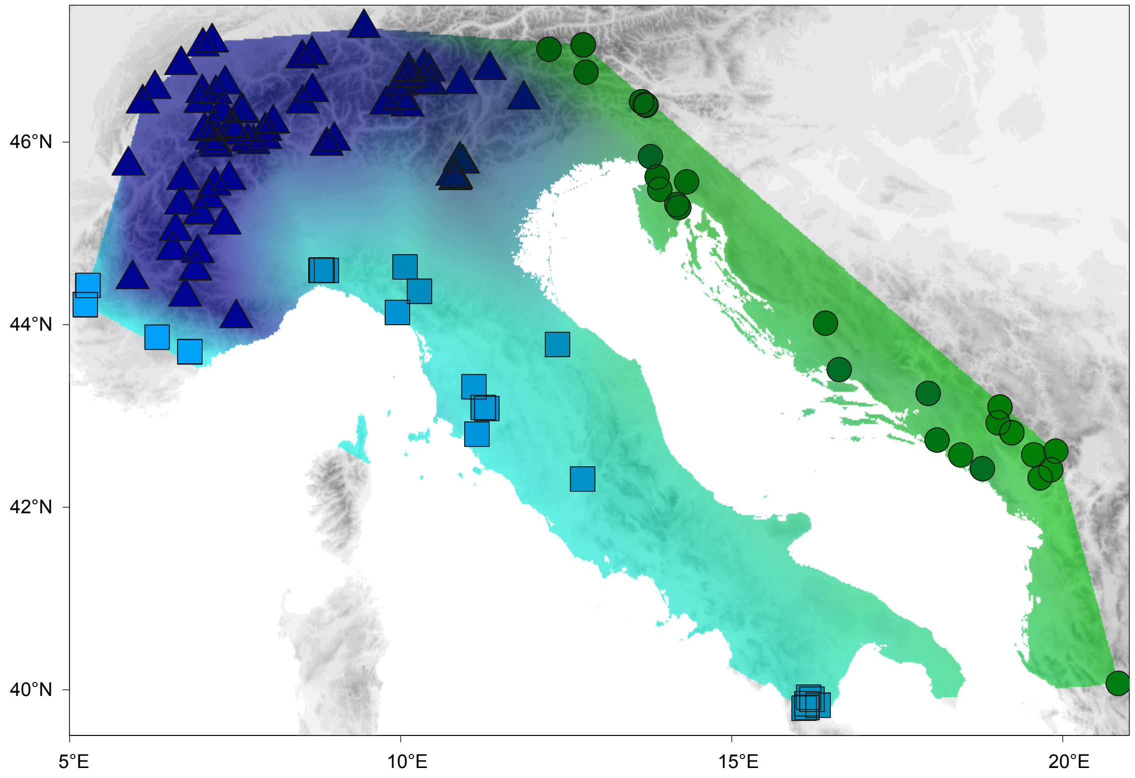


FIGURE S4. Spatial (linear) interpolation of the first two principle components of whole-genome variation (full dataset). Here, the first and second principal components are represented as green and blue colour channels, and explain 15.8% and 4.7% of the variance in the data, respectively. Individuals are denoted by shapes defined according to identified genetic clusters.

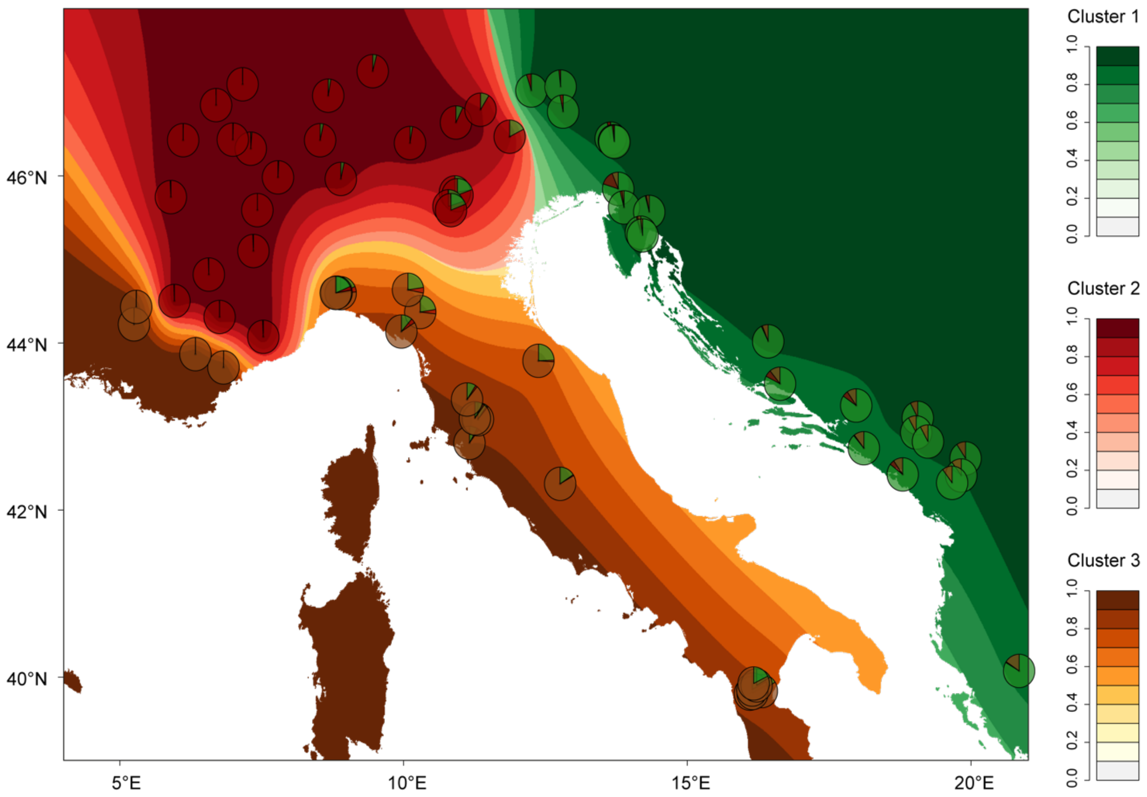


FIGURE S5. Population ancestry coefficients (K = 3) represented as pie charts and spatially interpolated (kriging) in geographic space. Contours may represent variations in ancestry coefficients (due to admixture) or alternatively, reflect recent bottleneck events. The latter interpretation is supported by patterns of allele sharing inferred via chromosome painting in the cases of the Alpine and Apennine lineages.

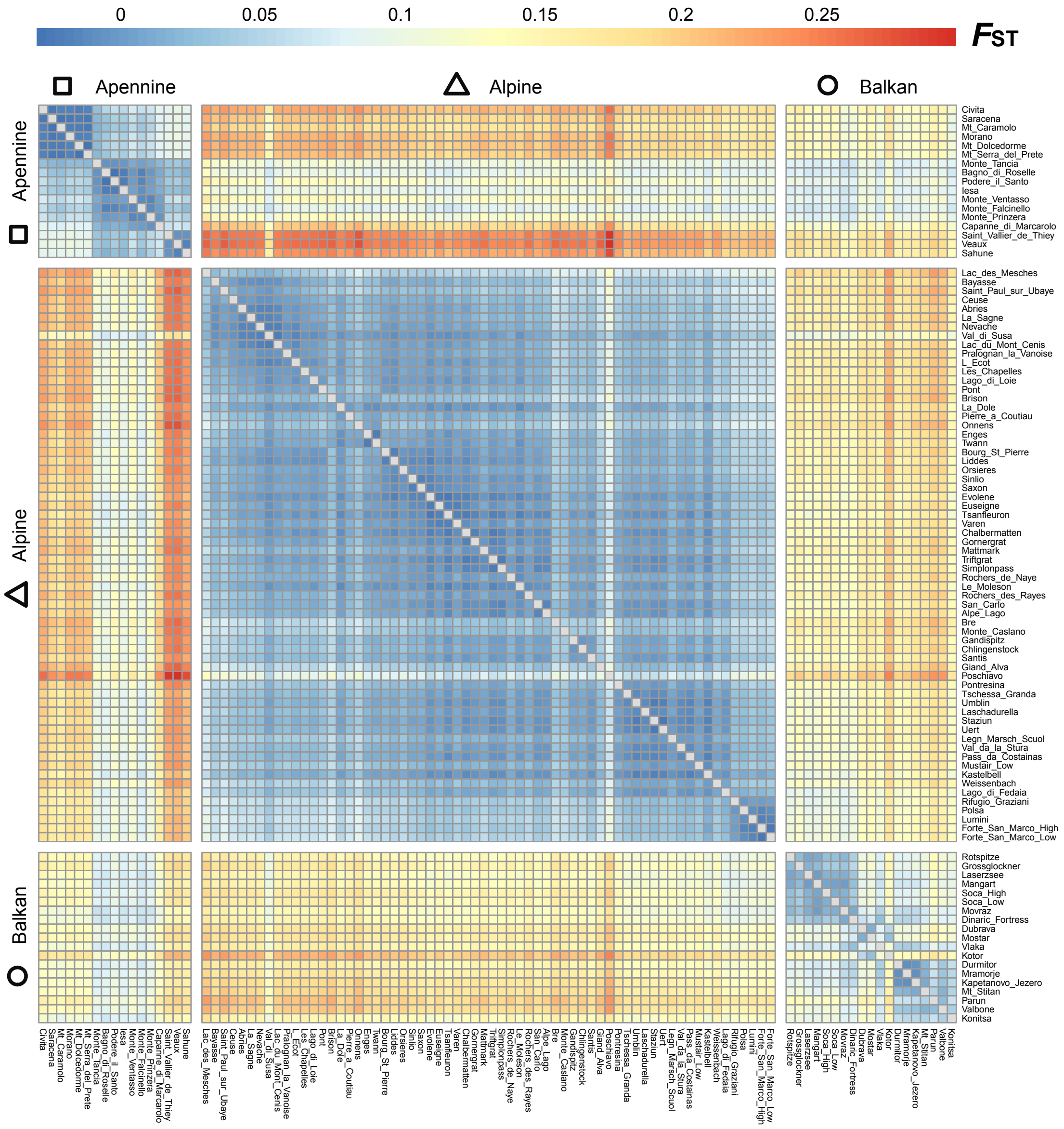


FIGURE S6. Heatmap of pairwise $F_{ST}$ between all populations. Populations are ordered (from left to right): Apennine lineage (south to north-west), Alpine linage (south-west to north-east), and Balkan lineage (north to south).

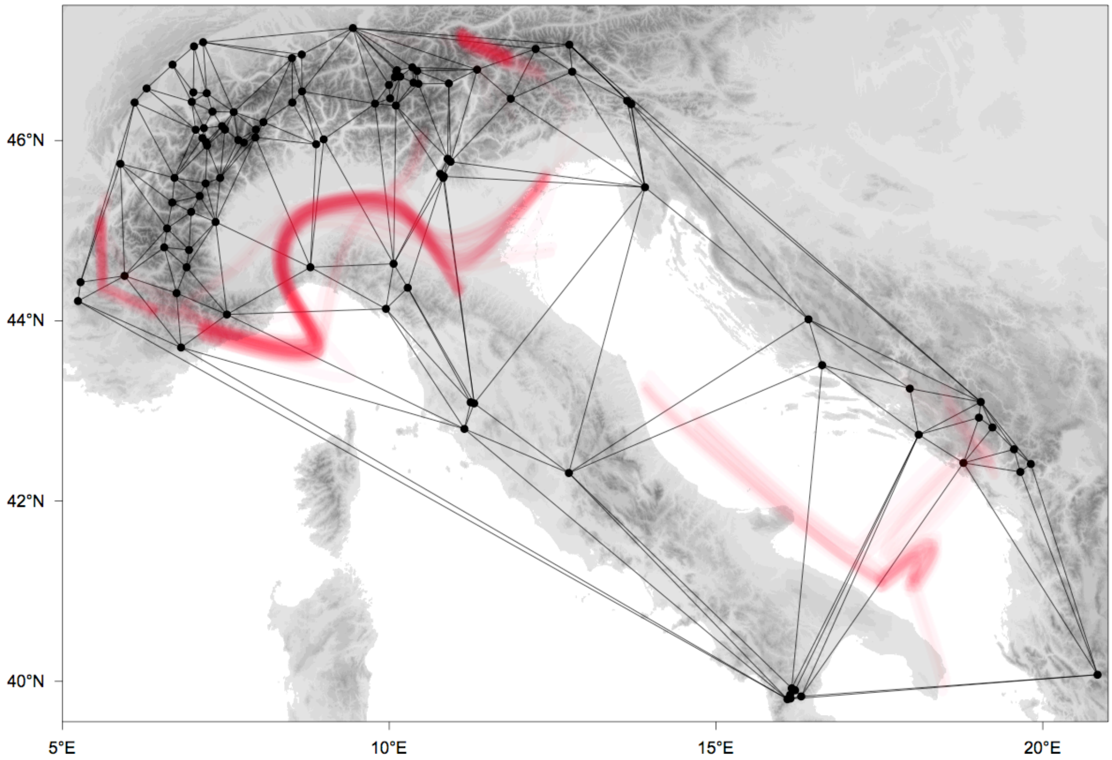


FIGURE S7. Genetic barriers (red) as identified in geographic space by Monmonier's algorithm applied on the $F_{ST}$ distance matrix. Populations are represented as vertices of a valuated graph constructed via Delauney triangulation, with edges reflecting population pairwise $F_{ST}$ (not scaled). The heatmap reflects support from 100 bootstrap runs.

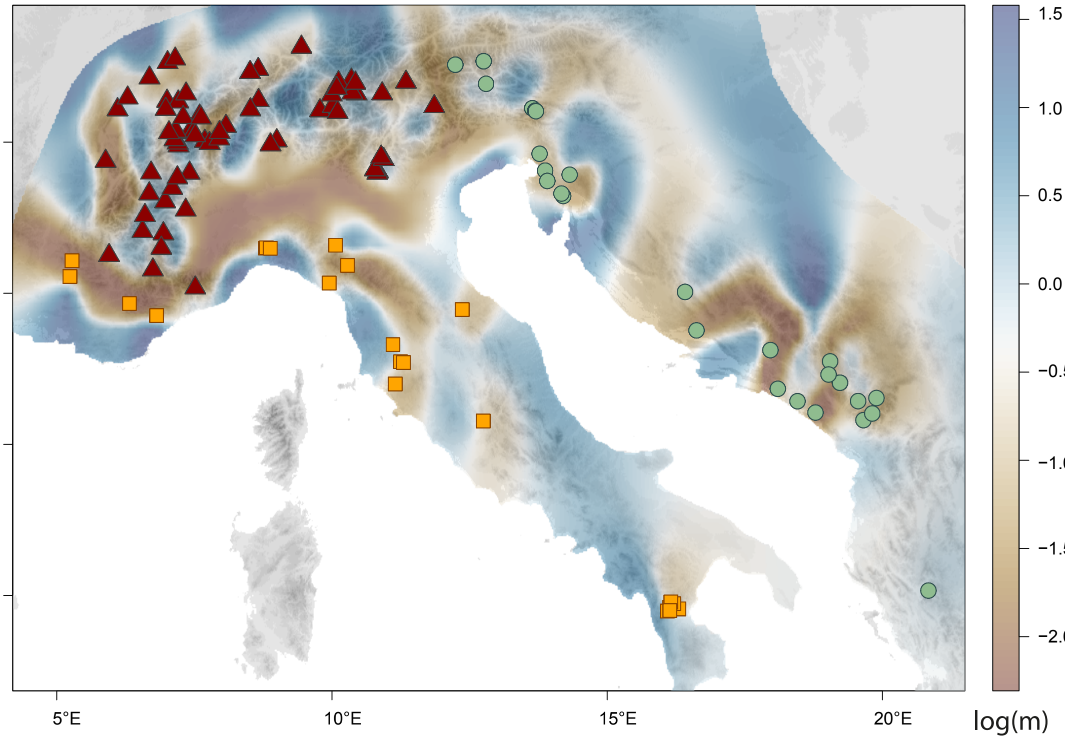


FIGURE S8. Map of effective migration rates (*m*), as inferred from *EEMS*. This map draws attention to where differentiation is elevated or depressed relative to expectations from geographic distance. Geographic locations of all samples are represented as coloured shapes, which are defined according to their genetic clusters.

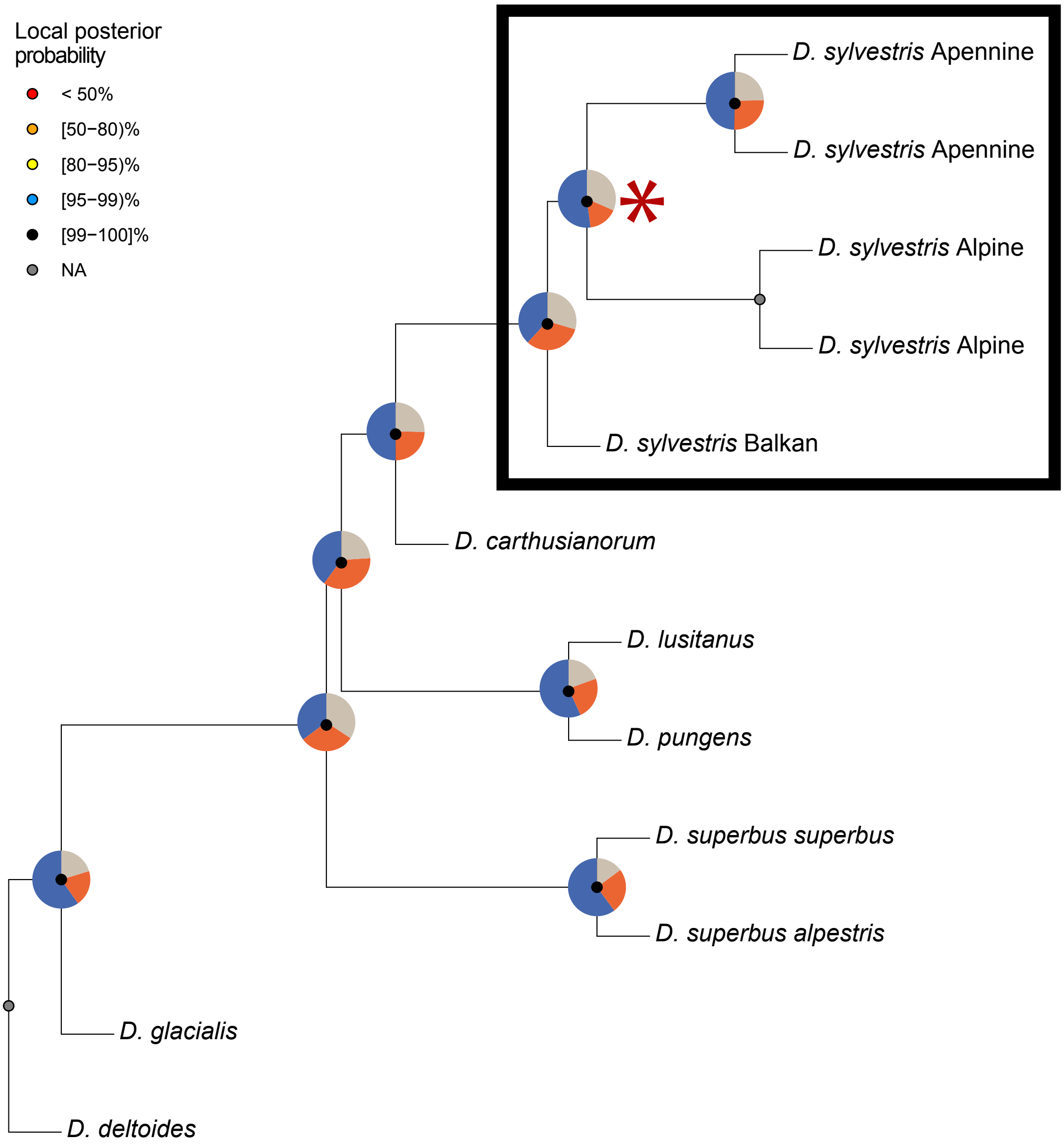


FIGURE S9. Distance-based phylogeny inferred using *ASTRAL-III* on 26,173 10-kb window (gene) trees. Taxon topology for *D. sylvestris* is shown in the black rectangle. The node defining the relationship between the three identified lineages of *D. sylvestris* is indicated with the red asterisk. Pie charts on nodes denote the fraction of gene trees that are consistent with the shown topology (blue), and with the first (orange) and second (grey) alternative topologies. Local posterior probabilities are shown as coloured circles on nodes. An alternate approach, based on concatenated whole-genome sequences, yielded an identical tree, with bootstrap values of 98-100% across all nodes, and is thus not shown.

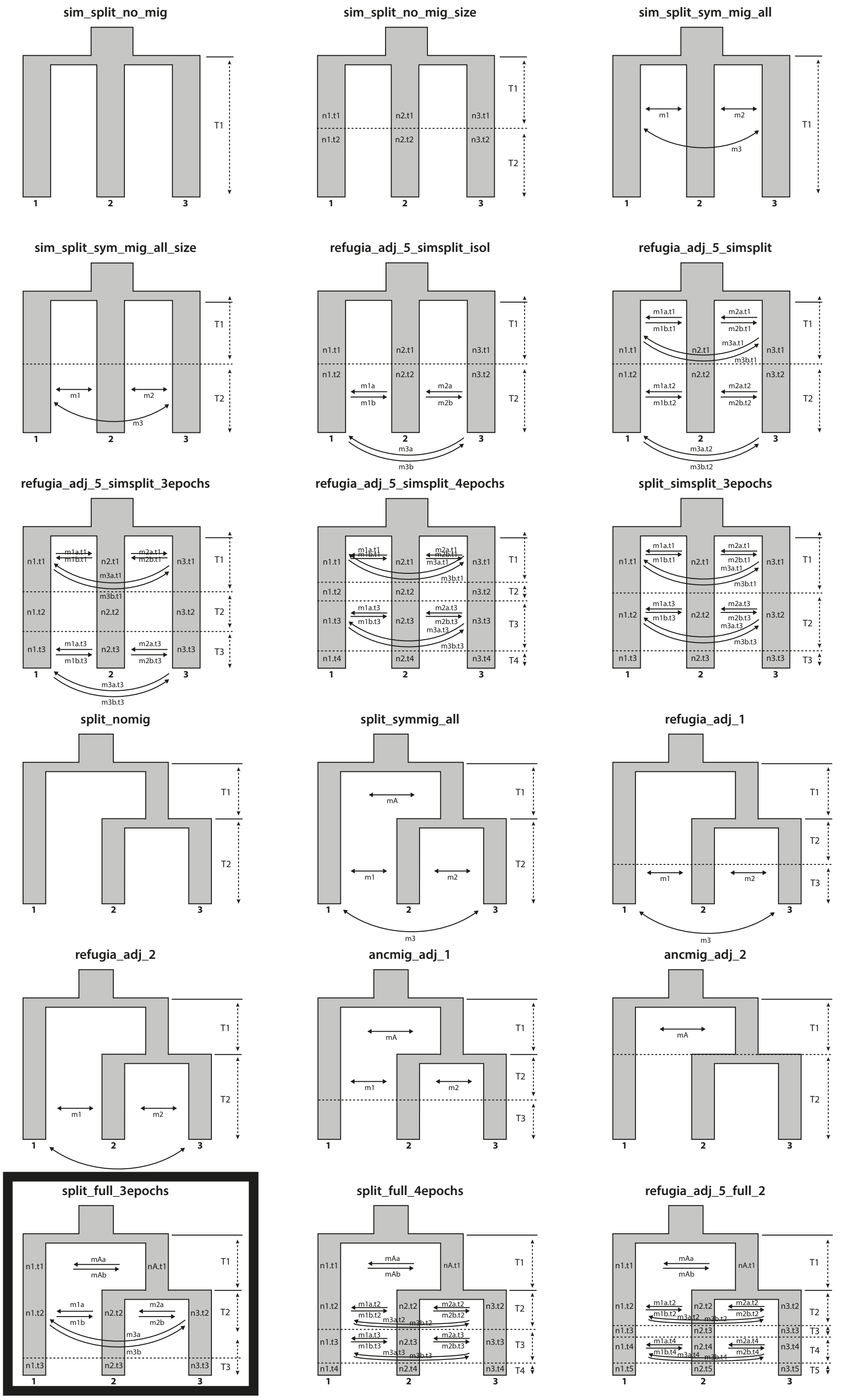


FIGURE S10. Demographic models tested. *y*-axis reflects time, with discrete time epochs denoted by T*i* ; $1 < i < 5$. Arrows represent migration, which may be unidirectional (one-headed arrow) or bidirectional (symmetric: single two-headed arrow; or asymmetric: two, opposing one-headed arrows). Population size change is allowed in certain models, and is denoted by n*j*.t*i*, where *j* = 1, 2, 3 refers to the Balkan, Apennine and Alpine lineages respectively. First and bottom three rows represent simultaneous and sequential trichotomous split topologies, respectively. The best model is indicated by black borders.

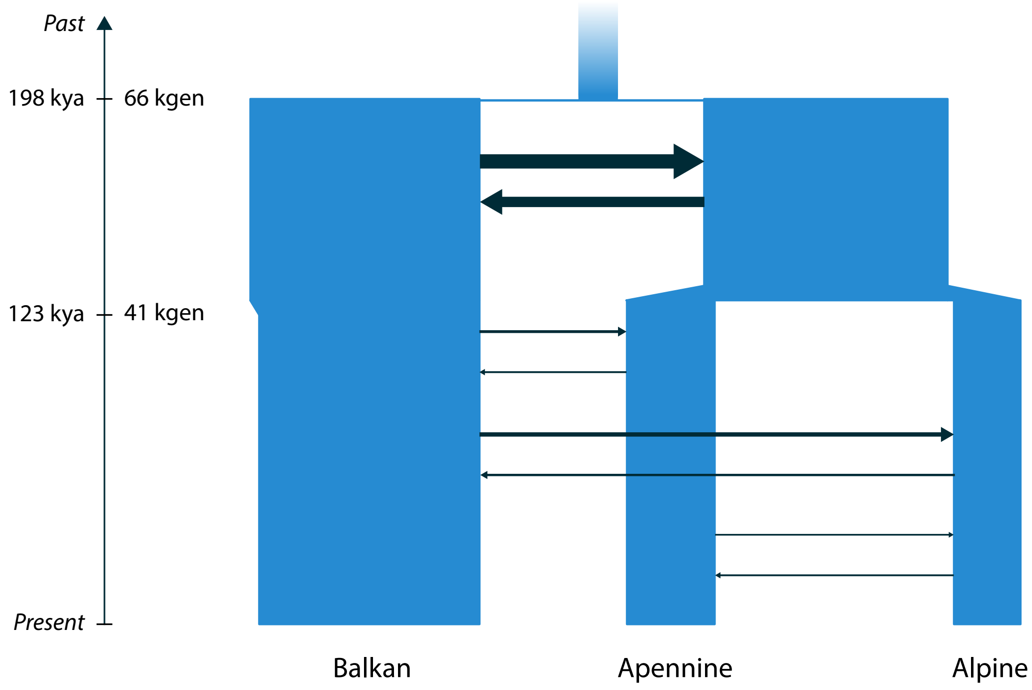


FIGURE S11. Best-fit demographic model estimates an initial split between lineages during the Penultimate Glacial Period ca. 198 kya (95% CI: 178-217 kya) followed by a subsequent split ca. 123 kya (95% CI: 114-132 kya). Here, time is shown both in units of generation time (thousands of generations, *kgen*) and absolute time (thousands of years, *kya*), with the latter assuming a generation time for *D. sylvestris* of 3 years. The last time epoch, reflective of the bottleneck-like effect of contemporary sampling of local populations, is not shown. Width of arrows and population blocks are proportional to estimated migration rates and effective population sizes, respectively.

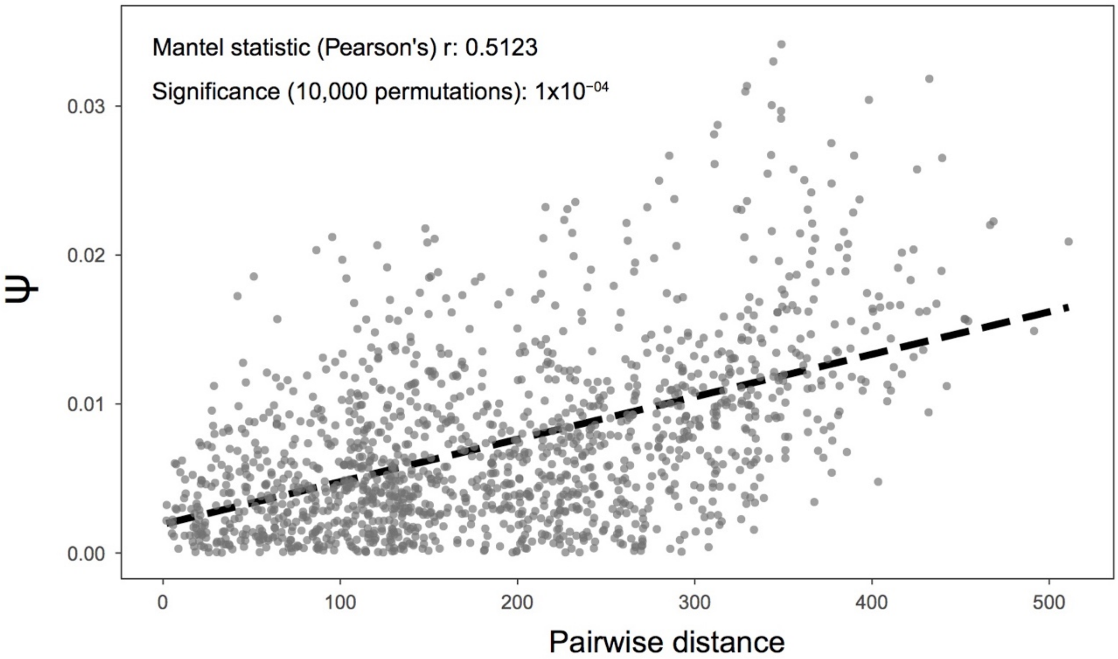


FIGURE S12. A significant, positive correlation of the directionality index $\psi$ with population pairwise geographic distance supports a spatial expansion of the Alpine lineage.

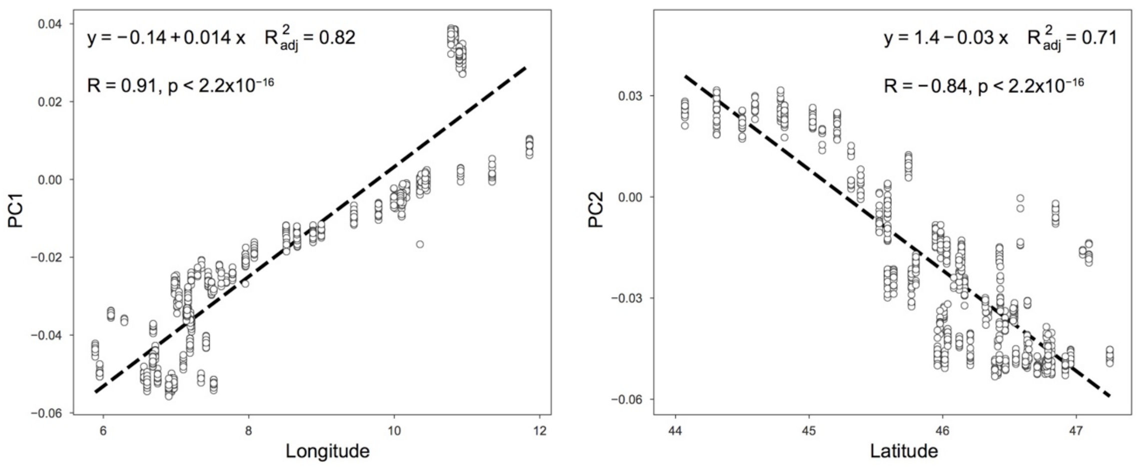


**FIGURE S13.** Correlation of longitude and latitude with the first two principle components of the PCA. The major axes of variation in genetic structure (PCs 1 and 2), under the default PCA rotation, were found to be maximally in line with the geographical axes, assessed via Procrustes rotational analyses.

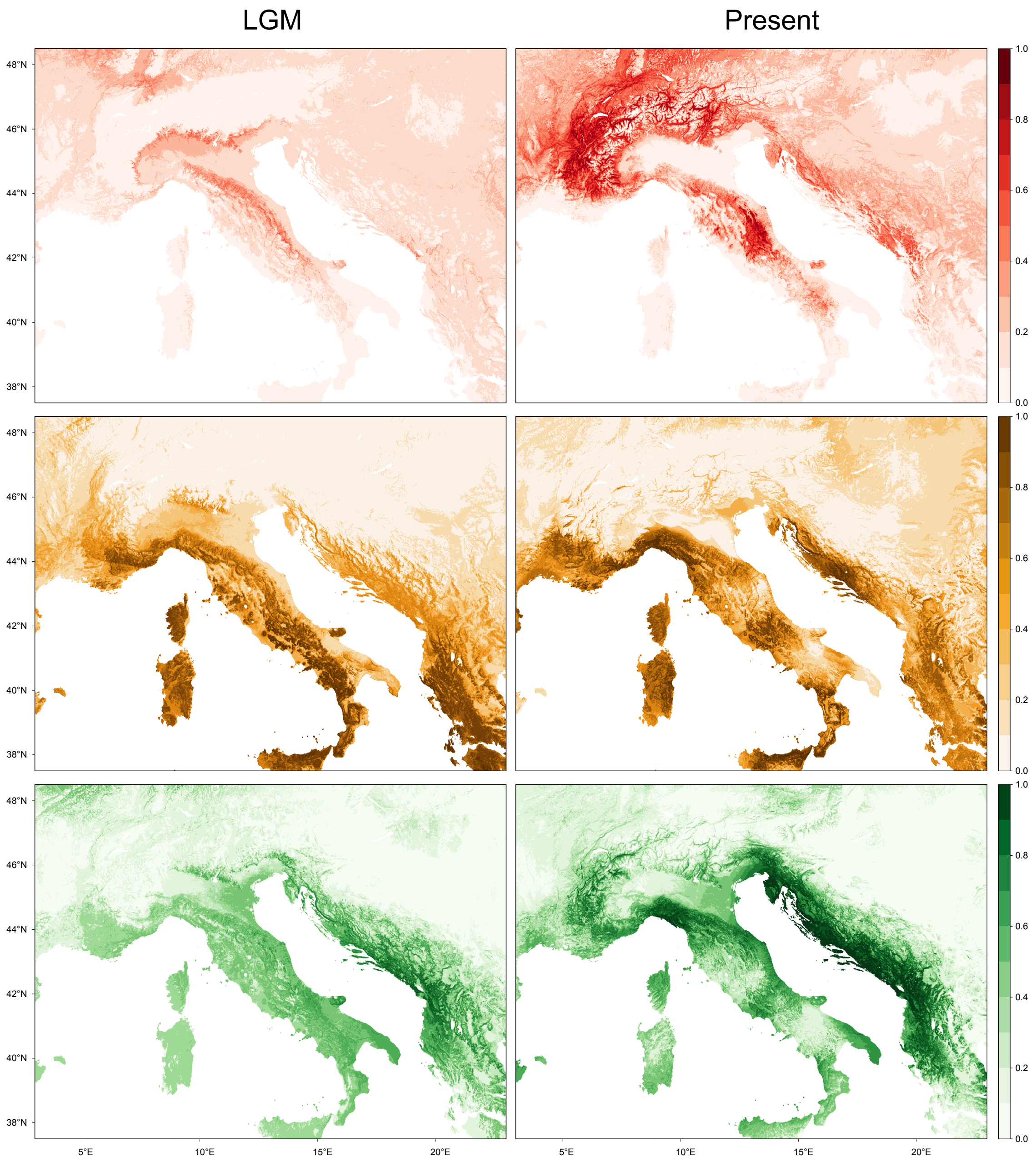


FIGURE S14. Predicted distribution models for the Alpine (red-top row), Apennine (yellow-middle row) and Balkan (green-bottom row) lineages, during the last glacial maximum LGM (left column) and present-day (right column). Darker hues reflect higher probabilities of presence.

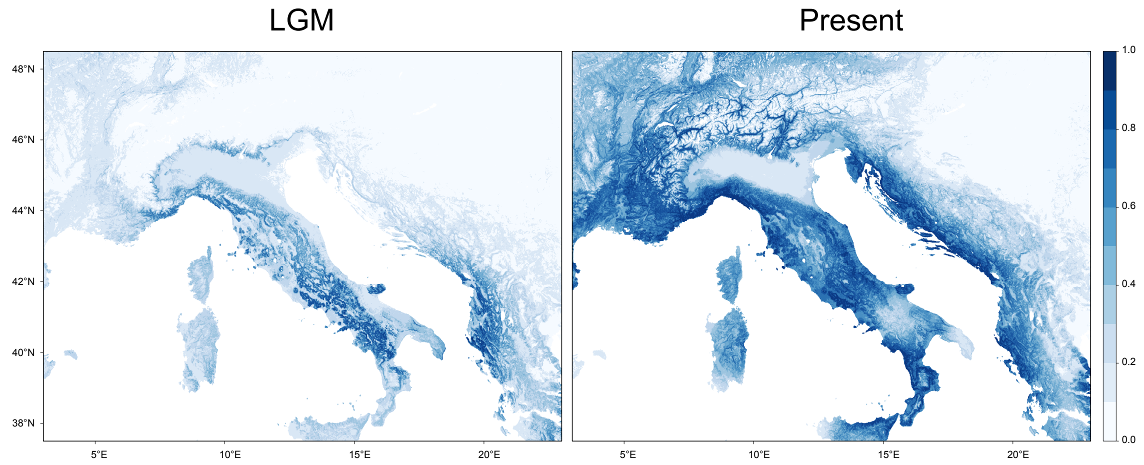


**FIGURE S15.** Predicted distribution models for the pooled species, during the last glacial maximum LGM (left column) and present-day (right column). Darker blue hues reflect higher probabilities of presence.

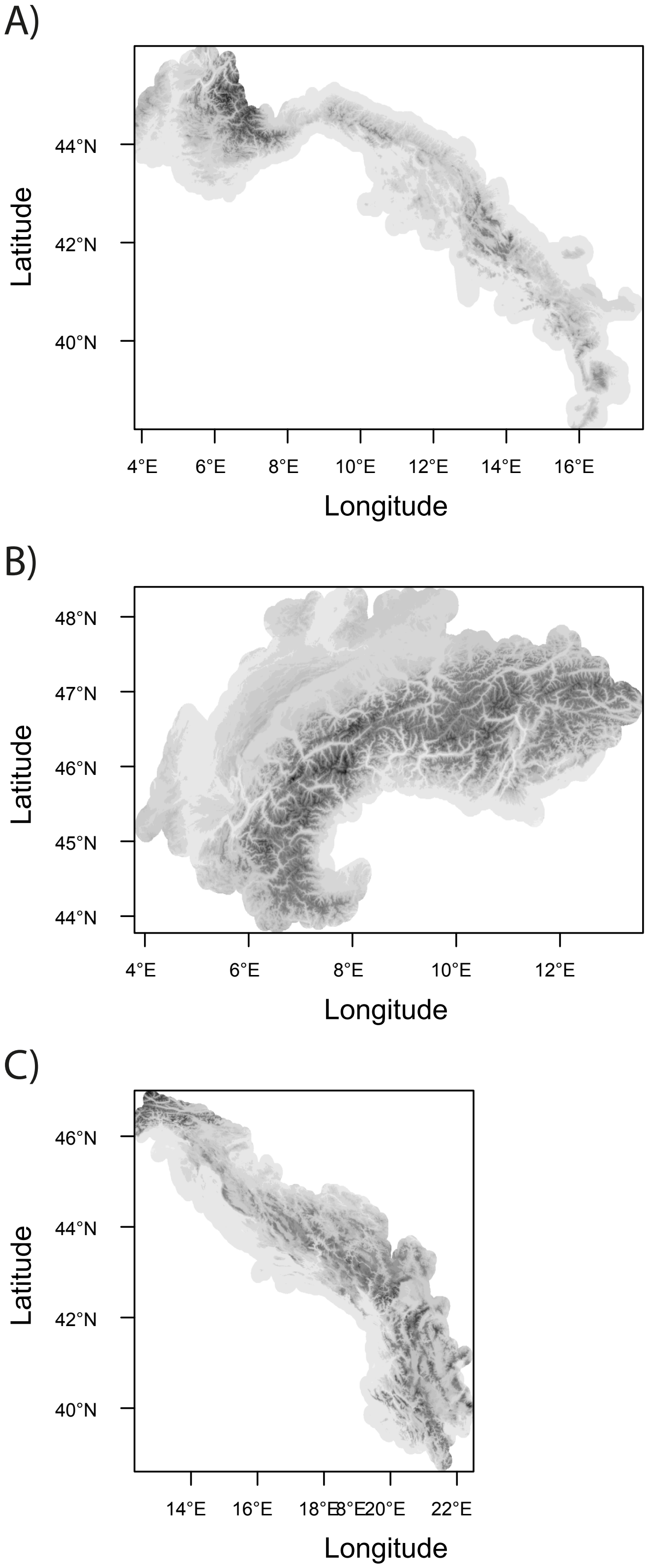


FIGURE S16. The geographic regions in which the environmental data was calculated, for the analyses of shifts in environmental space (**Fig. 4B**), for A) the Apennines, B) the Alps and C) the Balkans.

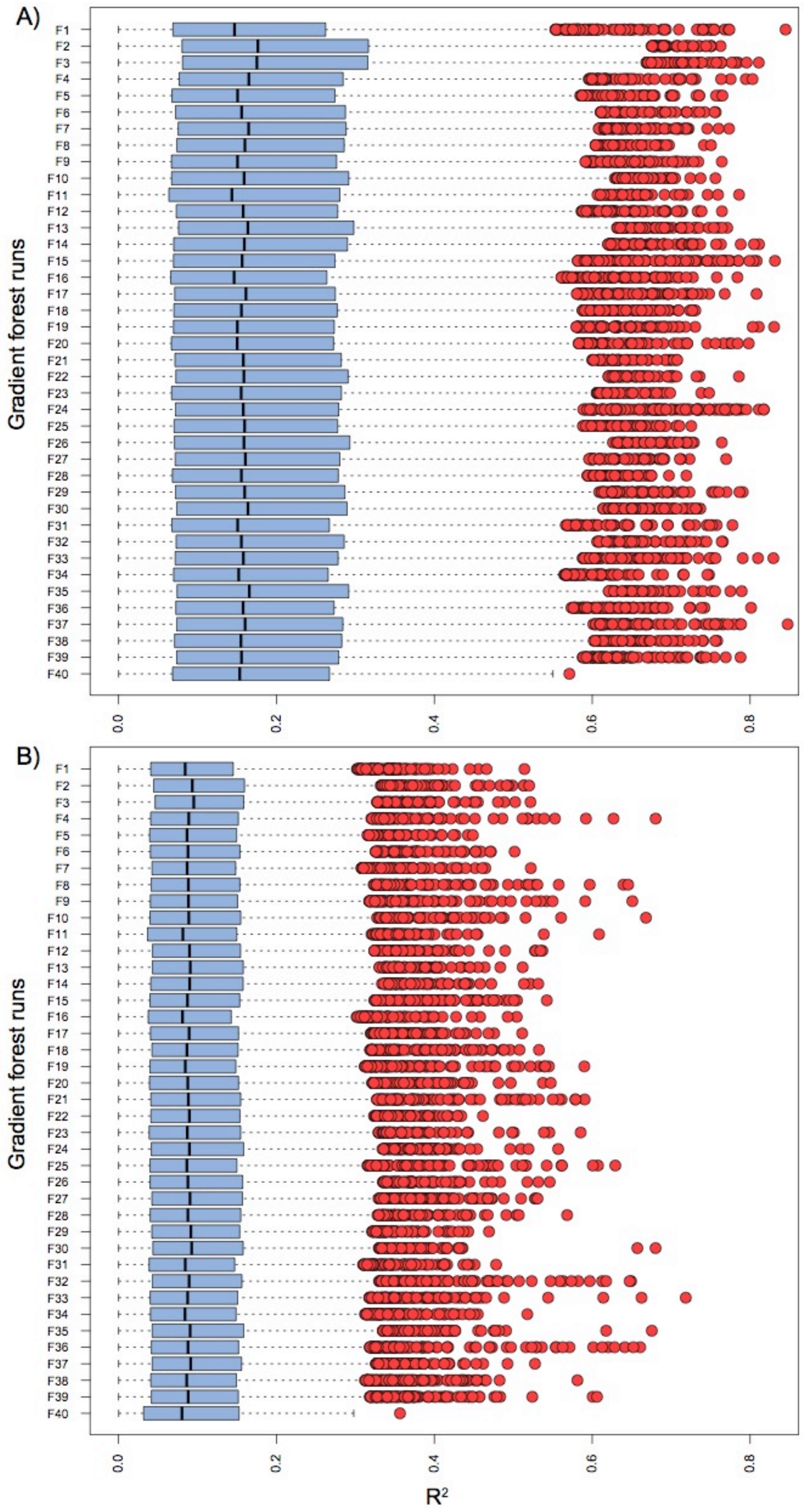


FIGURE S17. Box plots showing the variance in single nucleotide polymorphism (SNP) allele frequencies ($R^2$) explained by **A)** environmental + geographic predictor variables (12 variables total) and **B)** environmental predictor variables only (10 variables), under our gradient forest model. Due to the large number of SNPs, gradient forest was performed in partitions of 10,000 SNPs (390,262 SNPs total), resulting in 40 partitions, each of which is represented by a box plot here. Partitions were then combined into a single dataset.

### S3 SUPPLEMENTARY TABLES

| Population | Species | Latitude | Longitude | Elevation (m) | Date sampled | Country | Cluster | # individuals | PCA | Genetic distance | Admixture | badMIXTURE | Diversity | ψ | Demography | GF |
|---|---|---|---|---|---|---|---|---|---|---|---|---|---|---|---|---|
| Abries | *D. sylvestris* Wulfens | 44.786 | 6.937 | 1670 | 23.08.2016 | FR | Alpine | 15 | X | X | | | X | X | | X |
| Alpe_Lago | *D. sylvestris* Wulfens | 46.547 | 8.667 | 2150 | 16.07.2016 | CH | Alpine | 15 | X | X | | | X | X | | X |
| Bagno_di_Roselle | *D. sylvestris longicaulis* | 42.8 | 11.15 | 120 | 22.05.2011 | IT | Apennine | 5 | (X) | X | X | | X | | | |
| Bayasse | *D. sylvestris* Wulfens | 44.308 | 6.746 | 1868 | 25.08.2016 | FR | Alpine | 20 | (X) | X | X | X | X | X | | X |
| Bourg_St_Pierre | *D. sylvestris* Wulfens | 45.946 | 7.212 | 1780 | 22.08.2016 | CH | Alpine | 20 | X | X | | | X | X | | X |
| Bre | *D. sylvestris* Wulfens | 46.015 | 8.997 | 843 | 27.06.2017 | CH | Alpine | 14 | X | X | | | X | X | | X |
| Brison | *D. sylvestris* Wulfens | 45.744 | 5.888 | 206 | 10.06.2017 | FR | Alpine | 20 | (X) | X | X | | X | X | | X |
| Capanne_di_Marcarolo1 | *D.sylvestris* Wulfens | 44.595 | 8.794 | 570 | 31.01.2018 | IT | Apennine | 4 | (X) | X | X | X | X | | | |
| Capanne_di_Marcarolo2 | *D.sylvestris* Wulfens | 44.601 | 8.809 | 530 | 31.01.2018 | IT | Apennine | 4 | (X) | X | X | X | X | | | |
| Carpegna | *D. sylvestris longicaulis* | 43.783 | 12.367 | 700-800 | 26.08.2011 | IT | Apennine | 5 | (X) | X | X | | X | | | |
| Ceuse | *D. sylvestris* Wulfens | 44.499 | 5.95 | 1803 | 07.06.2017 | FR | Alpine | 15 | (X) | X | X | | X | X | | X |
| Chalbermatten | *D. sylvestris* Wulfens | 46.008 | 7.69 | 2100 | 12.07.2016 | CH | Alpine | 10 | X | X | | | X | X | | |
| Challer | *D. sylvestris* Wulfens | 46.341 | 7.595 | 2062 | 27.08.2012 | CH | Alpine | 5 | X | X | | | X | | | |
| Chlingenstock | *D.sylvestris* Wulfens | 46.957 | 8.661 | 1850 | 21.08.2016 | CH | Alpine | 15 | (X) | X | X | | X | X | | X |
| Civita | *D. sylvestris longicaulis* | 39.829 | 16.304 | 604 | 21.07.2015 | IT | Apennine | 20 | (X) | X | X | | X | | X | |
| Dinaric_Fortress | *D. sylvestris* Wulfens | 44.017 | 16.415 | 706 | 21.07.2017 | CR | Balkan | 5 | (X) | X | X | | X | | | |
| Dolina_Glinscice | *D. sylvestris tergestinus* | 45.62 | 13.874 | 130 | 21.06.2008 | IT | Balkan | 4 | (X) | X | X | | X | | | |
| Dubrava | *D. sylvestris* Wulfens | 43.508 | 16.624 | 471 | 22.07.2017 | CR | Balkan | 5 | (X) | X | X | | X | | | |
| Durmitor | *D. sylvestris* Wulfens | 43.099 | 19.051 | 1925 | 09.07.2008 | MN | Balkan | 5 | (X) | X | X | | X | | | |
| Enges | *D. sylvestris* Wulfens | 47.048 | 7.013 | 673 | 28.07.2016 | CH | Alpine | 5 | X | X | | | X | | | |
| Euseigne | *D. sylvestris* Wulfens | 46.163 | 7.441 | 1020 | 24.08.2016 | CH | Alpine | 20 | X | X | | | X | X | | X |
| Evolene | *D. sylvestris* Wulfens | 46.123 | 7.487 | 1520 | 23.08.2016 | CH | Alpine | 20 | X | X | | | X | X | | X |
| Forte_San_Marco_High | *D. sylvestris* Wulfens | 45.603 | 10.836 | 500 | 16.07.2017 | IT | Alpine | 20 | X | X | | X | X | X | | X |
| Forte_San_Marco_Low | *D. sylvestris* Wulfens | 45.587 | 10.825 | 235 | 16.07.2017 | IT | Alpine | 20 | (X) | X | X | | X | X | | X |
| Fraconalto | *D. sylvestris* Wulfens | 44.593 | 8.88 | 720 | 31.01.2018 | IT | Apennine | 4 | (X) | X | X | X | X | | | |
| Gandispitz | *D. sylvestris* Wulfens | 46.917 | 8.513 | 2013 | 08.09.2016 | CH | Alpine | 13 | X | X | | | X | X | | |
| Giand_Alva | *D. sylvestris* Wulfens | 46.412 | 9.783 | 2430-2440 | 29.07.2016 | CH | Alpine | 14 | X | X | | | X | X | | X |
| Gornergrat | *D. sylvestris* Wulfens | 45.979 | 7.776 | 2740 | 13.07.2016 | CH | Alpine | 15 | (X) | X | X | | X | X | | X |
| Grossglockner | *D. sylvestris* Wulfens | 47.066 | 12.756 | 2104 | 18.08.2017 | AT | Balkan | 15 | (X) | X | X | X | X | | | |
| Hauderes | *D. sylvestris* Wulfens | 46.082 | 7.518 | 1595 | 23.08.2016 | CH | Alpine | 5 | X | | | | | | | |
| Iesa | *D. sylvestris longicaulis* | 43.095 | 11.246 | 340 | 21.06.2011 | IT | Apennine | 5 | (X) | X | X | | X | | | |
| Kapetanovo_Jezero | *D. sylvestris* Wulfens | 42.816 | 19.23 | 1741 | 25.07.2017 | MN | Balkan | 20 | (X) | X | X | | X | | X | |
| Kastelbell | *D. sylvestris* Wulfens | 46.636 | 10.912 | 690-770 | 08.08.2016 | IT | Alpine | 10 | (X) | X | X | | X | X | | |

| | | | | | | | | | | | | | | | | |
|---|---|---|---|---|---|---|---|---|---|---|---|---|---|---|---|---|
| **Komen** | *D. sylvestris tergestinus* | 45.842 | 13.772 | 294 | 20.06.2017 | SL | Balkan | 3 | (X) | X | X | | X | | | |
| **Konitsa** | *D. sylvestris* Wulfens | 40.07 | 20.838 | 900-950 | 20.07.2011 | GR | Balkan | 5 | (X) | X | X | | X | | | |
| **Kotor** | *D. sylvestris* Wulfens | 42.422 | 18.786 | 629.7 | 24.05.2017 | MN | Balkan | 5 | (X) | X | X | | X | | | |
| **L_Ecot** | *D. sylvestris* Wulfens | 45.386 | 7.1 | 2090 | 20.08.2016 | FR | Alpine | 10 | X | X | | | X | X | | |
| **La_Dole** | *D. sylvestris* Wulfens | 46.425 | 6.104 | 1456 | 29.07.2016 | CH | Alpine | 15 | (X) | X | X | | X | X | | X |
| **La_Sagne** | *D. sylvestris* Wulfens | 44.816 | 6.557 | 1016 | 09.06.2017 | FR | Alpine | 20 | (X) | X | X | X | X | X | | X |
| **Lac_des_Mesches** | *D. sylvestris* Wulfens | 44.07 | 7.517 | 1453 | 04.06.2017 | FR | Alpine | 15 | (X) | X | X | | X | X | | X |
| **Lac_du_Mont_Cenis** | *D. sylvestris* Wulfens | 45.209 | 6.97 | 1938 | 21.08.2016 | FR | Alpine | 20 | X | X | | | X | X | | X |
| **Lago_di_Fedaia** | *D. sylvestris* Wulfens | 46.466 | 11.859 | 2220 | 04.08.2016 | IT | Alpine | 15 | (X) | X | X | | X | X | | X |
| **Lago_di_Loie** | *D. sylvestris* Wulfens | 45.587 | 7.413 | 2365 | 29.09.2016 | IT | Alpine | 15 | (X) | X | X | | X | X | | X |
| **Laschadurella** | *D. sylvestris* Wulfens | 46.712 | 10.167 | 2340-2410 | 16.07.2016 | CH | Alpine | 10 | X | X | | | X | X | | |
| **Laserzsee** | *D. sylvestris* Wulfens | 46.767 | 12.8 | 2232 | 19.08.2017 | AT | Balkan | 5 | (X) | X | X | X | X | | | |
| **Le_Moleson** | *D. sylvestris* Wulfens | 46.537 | 7.004 | 1820 | 25.07.2016 | CH | Alpine | 15 | X | X | | | X | X | | X |
| **Legn_Marsch_Scuol** | *D. sylvestris* Wulfens | 46.815 | 10.353 | 1080-1280 | 23.06.2016 | CH | Alpine | 15 | X | X | | | X | X | | X |
| **Les_Chapelles** | *D. sylvestris* Wulfens | 45.587 | 6.714 | 1432 | 18.08.2016 | FR | Alpine | 15 | X | X | | | X | X | | X |
| **Liddes** | *D. sylvestris* Wulfens | 45.984 | 7.197 | 1530 | 22.08.2016 | CH | Alpine | 20 | X | X | | | X | X | | X |
| **Lumini** | *D. sylvestris* Wulfens | 45.632 | 10.78 | 1051 | 17.07.2017 | IT | Alpine | 20 | (X) | X | X | X | X | X | | X |
| **Mangart** | *D. sylvestris* Wulfens | 46.444 | 13.64 | 2053 | 08.08.2016 | SL | Balkan | 20 | (X) | X | X | X | X | | | |
| **Mattmark** | *D. sylvestris* Wulfens | 46.037 | 7.953 | 2340 | 04.08.2016 | CH | Alpine | 15 | X | X | | | X | X | | X |
| **Mont_Chiran** | *D. sylvestris longicaulis* | 43.863 | 6.321 | 1705 | 06.06.2017 | FR | Apennine | 4 | (X) | X | X | X | X | | | |
| **Monte_Caslano** | *D. sylvestris* Wulfens | 45.961 | 8.883 | 332 | 26.06.2017 | CH | Alpine | 15 | (X) | X | X | | X | X | | X |
| **Monte_Falcinello** | *D. sylvestris longicaulis* | 44.133 | 9.95 | 140 | 2011 | IT | Apennine | 5 | (X) | X | X | | X | | | |
| **Monte_Prinzera** | *D. sylvestris longicaulis* | 44.633 | 10.067 | 590-600 | 08.04.2011 | IT | Apennine | 5 | (X) | X | X | X | X | | | |
| **Monte_Tancia** | *D. sylvestris longicaulis* | 42.311 | 12.749 | 900 | 13.05.2011 | IT | Apennine | 5 | (X) | X | X | | X | | | |
| **Monte_Ventasso** | *D. sylvestris longicaulis* | 44.367 | 10.283 | 1680 | 2011 | IT | Apennine | 5 | (X) | X | X | X | X | | | |
| **Morano** | *D. sylvestris longicaulis* | 39.847 | 16.134 | 585 | 22.07.2015 | IT | Apennine | 5 | (X) | X | X | | X | | | |
| **Mostar** | *D. sylvestris* Wulfens | 43.248 | 17.966 | 827 | 26.07.2017 | BO | Balkan | 5 | (X) | X | X | | X | | | |
| **Movraz** | *D. sylvestris tergestinus* | 45.483 | 13.913 | 410 | 27.06.2017 | SL | Balkan | 5 | X | X | | | X | | | |
| **Mramorje** | *D. sylvestris* Wulfens | 42.925 | 19.023 | 1610 | 08.07.2008 | MN | Balkan | 5 | (X) | X | X | | X | | | |
| **Mt_Caramolo** | *Dianthus brachycalyx* | 39.798 | 16.092 | 1824 | 04.08.2015 | IT | Apennine | 5 | (X) | X | X | | X | | | |
| **Mt_Dolcedorme** | *Dianthus brachycalyx* | 39.897 | 16.21 | 2134 | 13.08.2015 | IT | Apennine | 5 | (X) | X | X | | X | | | |
| **Mt_Serra_del_Prete** | *Dianthus brachycalyx* | 39.92 | 16.158 | 2181 | 06.08.2015 | IT | Apennine | 5 | (X) | X | X | | X | | | |
| **Mt_Stitan** | *D. sylvestris bertisceus* | 42.574 | 19.56 | 2000 | 08.07.2017 | MN | Balkan | 5 | X | X | | | X | | | |
| **Mustair_Low** | *D. sylvestris* Wulfens | 46.632 | 10.447 | 1300 | 03.08.2016 | CH | Alpine | 15 | X | X | | | X | X | | X |
| **Nevache** | *D. sylvestris* Wulfens | 45.027 | 6.602 | 1871 | 22.08.2016 | FR | Alpine | 20 | X | X | | X | X | X | | X |
| **Onnens** | *D. sylvestris* Wulfens | 46.845 | 6.685 | 488 | 28.07.2016 | CH | Alpine | 14 | (X) | X | X | | X | X | | X |
| **Orjen** | *D. sylvestris nodosus* | 42.572 | 18.459 | 910 | 11.06.2017 | BO | Balkan | 5 | X | X | | | X | | | |
| **Orsieres** | *D. sylvestris* Wulfens | 46.03 | 7.141 | 990 | 23.08.2016 | CH | Alpine | 20 | X | X | | | X | X | | X |
| **Parun** | *D. sylvestris* Wulfens. | 42.322 | 19.657 | 1300 | 17.07.2008 | AL | Balkan | 5 | (X) | X | X | | X | | | |
| **Pass_da_Costainas** | *D. sylvestris* Wulfens | 46.644 | 10.372 | 2260-2340 | 24.07.2016 | CH | Alpine | 5 | X | X | | | X | | | |
| **Pierre_a_Coutiau** | *D. sylvestris* Wulfens | 46.581 | 6.288 | 1608 | 29.07.2016 | CH | Alpine | 5 | X | X | | | X | | | |

| | | | | | | | | | | | | | | | | |
|---|---|---|---|---|---|---|---|---|---|---|---|---|---|---|---|---|
| **Pievescola** | *D. sylvestris. longicaulis* | 43.32 | 11.108 | 230 | 2011 | IT | Apennine | 5 | (X) | X | X | | X | | | |
| **Podere_il_Santo** | *D. sylvestris longicaulis* | 43.083 | 11.3 | 275-350 | 26.06.2011 | IT | Apennine | 5 | (X) | X | X | | X | | | |
| **Polsa** | *D. sylvestris* Wulfens | 45.766 | 10.939 | 1350 | 15.07.2017 | IT | Alpine | 20 | (X) | X | X | X | X | X | X | X |
| **Pont** | *D. sylvestris* Wulfens | 45.525 | 7.191 | 2130 | 28.09.2016 | IT | Alpine | 14 | X | X | | | X | X | | X |
| **Pontresina** | *D. sylvestris* Wulfens | 46.469 | 10.014 | 2520-2545 | 28.07.2016 | CH | Alpine | 10 | X | X | | | X | X | | |
| **Poschiavo** | *D. sylvestris* Wulfens | 46.39 | 10.102 | 1874 | 28.09.2017 | CH | Alpine | 7 | (X) | X | X | | X | | | |
| **Pralognan_la_Vanoise** | *D. sylvestris* Wulfens | 45.314 | 6.68 | 1986 | 19.08.2016 | FR | Alpine | 15 | X | X | | | X | X | | X |
| **Rifugio_Graziani** | *D. sylvestris* Wulfens | 45.799 | 10.894 | 1566 | 15.07.2017 | IT | Alpine | 20 | (X) | X | X | X | X | X | | X |
| **Rochers_de_Naye** | *D. sylvestris* Wulfens | 46.431 | 6.98 | 1975 | 25.07.2016 | CH | Alpine | 15 | (X) | X | X | | X | X | | X |
| **Rochers_des_Rayes** | *D. sylvestris* Wulfens | 46.527 | 7.208 | 1790 | 24.07.2016 | CH | Alpine | 10 | X | X | | | X | | | |
| **Rotspitze** | *D. sylvestris* Wulfens | 47.019 | 12.241 | 2070 | 17.08.2017 | AT | Balkan | 15 | (X) | X | X | X | X | | | |
| **Sahune** | *D. sylvestris longicaulis* | 44.429 | 5.277 | 428 | 07.06.2017 | FR | Apennine | 15 | (X) | X | X | | X | | | |
| **Saint_Paul_sur_Ubaye** | *D. sylvestris* Wulfens | 44.595 | 6.9 | 2343 | 24.08.2016 | FR | Alpine | 20 | X | X | | X | X | X | | X |
| **Saint_Vallier_de_Thiey** | *D. sylvestris longicaulis* | 43.704 | 6.815 | 900 | 05.06.2017 | FR | Apennine | 15 | (X) | X | X | | X | | | |
| **San_Carlo** | *D. sylvestris* Wulfens | 46.424 | 8.518 | 1420 | 01.07.2016 | CH | Alpine | 10 | (X) | X | X | | X | X | | |
| **Santis** | *D. sylvestris* Wulfens | 47.251 | 9.445 | 1512 | 27.09.2017 | CH | Alpine | 15 | (X) | X | X | | X | X | | X |
| **Saracena** | *D. sylvestris longicaulis* | 39.806 | 16.14 | 909 | 30.07.2015 | IT | Apennine | 5 | (X) | X | X | | X | | | |
| **Saxon** | *D. sylvestris* Wulfens | 46.14 | 7.164 | 570 | 28.06.2012 | CH | Alpine | 20 | X | X | | | X | X | | X |
| **Schwarzenmatt** | *D. sylvestris* Wulfens | 46.634 | 7.349 | 1220 | 23.07.2016 | CH | Alpine | 5 | X | X | | | X | | | |
| **Simplonpass** | *D. sylvestris* Wulfens | 46.206 | 8.074 | 2280 | 06.08.2016 | CH | Alpine | 15 | X | X | | | X | X | | X |
| **Sinlio** | *D. sylvestris* Wulfens | 46.122 | 7.039 | 900 | 17.08.2016 | CH | Alpine | 5 | X | X | | | X | | | |
| **Sneznik** | *D. sylvestris* Wulfens | 45.565 | 14.318 | 1000 | 18.07.2017 | SL | Balkan | 3 | (X) | X | X | | X | | | |
| **Soca_High** | *D. sylvestris* Wulfens | 46.412 | 13.694 | 1632 | 09.08.2016 | SL | Balkan | 5 | (X) | X | X | X | X | | | |
| **Soca_Low** | *D. sylvestris* Wulfens | 46.405 | 13.705 | 1036 | 09.08.2016 | SL | Balkan | 5 | (X) | X | X | X | X | | | |
| **Staziun** | *D. sylvestris* Wulfens | 46.767 | 10.108 | 1440-1500 | 23.06.2016 | CH | Alpine | 10 | X | X | | | X | X | | |
| **Triftgrat** | *D. sylvestris* Wulfens | 46.124 | 7.958 | 2500 | 05.08.2016 | CH | Alpine | 15 | X | X | | | X | X | | X |
| **Tsanfleuron** | *D. sylvestris* Wulfens | 46.323 | 7.296 | 2110 | | CH | Alpine | 5 | (X) | X | X | | X | | | |
| **Tschessa_Granda** | *D. sylvestris* Wulfens | 46.619 | 9.995 | 1700 | 09.07.2017 | CH | Alpine | 10 | X | X | | | X | X | | |
| **Twann** | *D. sylvestris* Wulfens | 47.094 | 7.152 | 490 | 28.07.2016 | CH | Alpine | 10 | (X) | X | X | | X | X | | |
| **Ucka** | *D. sylvestris* Wulfens | 45.284 | 14.202 | 1400 | 20.06.2017 | CR | Balkan | 3 | (X) | X | X | | X | | | |
| **Uert** | *D. sylvestris* Wulfens | 46.782 | 10.119 | 2090-2115 | 07.07.2016 | CH | Alpine | 10 | X | X | | | X | X | | |
| **Umblin** | *D. sylvestris* Wulfens | 46.707 | 10.087 | 1480-1535 | 23.06.2016 | CH | Alpine | 10 | X | X | | | X | | | |
| **Val_da_la_Stura** | *D. sylvestris* Wulfens | 46.775 | 10.426 | 2160-2365 | 01.07.2016 | CH | Alpine | 15 | X | X | | | X | X | | X |
| **Val_di_Susa** | *D. sylvestris* Wulfens | 45.098 | 7.343 | 880 | 2011 | IT | Alpine | 7 | (X) | X | X | | X | | | |
| **Valbone** | *D. sylvestris* Wulfens | 42.41 | 19.817 | 1530-1800 | 29.07.2017 | AL | Balkan | 5 | (X) | X | X | | X | | | |
| **Varen** | *D. sylvestris* Wulfens | 46.318 | 7.621 | 768 | 29.06.2012 | CH | Alpine | 5 | X | X | | | X | | | |
| **Veaux** | *D. sylvestris longicaulis* | 44.22 | 5.238 | 569 | 06.06.2017 | FR | Apennine | 20 | (X) | X | X | X | X | | | |
| **Vela_Draga** | *D. sylvestris* Wulfens | 45.317 | 14.17 | 470 | 20.06.2017 | CR | Balkan | 3 | (X) | X | X | | X | | | |
| **Visitor** | *D. sylvestris bertisceus* | 42.615 | 19.893 | 1850 | 08.07.2008 | HU | Balkan | 3 | (X) | X | X | | X | | | |
| **Vlaka** | *D. sylvestris* Wulfens | 42.736 | 18.101 | 525 | 23.07.2017 | BO | Balkan | 5 | (X) | X | X | | X | | | |
| **Weissenbach** | *D. sylvestris* Wulfens | 46.789 | 11.345 | 1774 | 14.07.2017 | IT | Alpine | 15 | (X) | X | X | | X | X | | X |

**Table S1: Sampling of wild populations.** Population name, taxonomic denomination, geographic coordinates (WGS 84), elevation (meters above sea level), date sampled, country (ISO 3166-1 alpha-2 codes), and number of sequenced individuals per population are reported. Note that taxonomic names adhere to established epithets used in floristic treatments. Revision of the taxonomy of *D. sylvestris* s.l. is ongoing and will account for the results of this study. The populations used in the following population genetic analyses are reported: PCA (all individuals from all populations; alternative run on 125 individuals per meta-population indicated in parentheses), genetic distance (5 individuals per population), admixture (125 individuals per meta-population), *badMIXTURE* (22 and 65 individuals per meta-population, for the Apennine-Balkan and Alpine-Balkan analyses respectively), diversity (*EEMS*; 5 individuals per population), $\psi$ (10 individuals per population), demography (20 individuals per population), and gradient forest (GF; 14 individuals per population).

| Model | log-likelihood | AIC |
|---|---|---|
| split_full_3epochs | -901193 | 1802424 |
| split_full_4epochs | -903122 | 1806303 |
| split_simsplit_3epochs | -907979 | 1816007 |
| refugia_adj_5_simsplit_4epochs | -923895 | 1847846 |
| refugia_adj_5_simsplit_isol | -946667 | 1893361 |
| refugia_adj_5_simsplit_3epochs | -952499 | 1905045 |
| sim_split_sym_mig_all_size | -954060 | 1908144 |
| split_symmig_all | -958882 | 1917784 |
| sim_split_sym_mig_all | -959422 | 1918860 |
| refugia_adj_2 | -960327 | 1920673 |
| refugia_adj_1 | -960596 | 1921213 |
| refugia_adj_5_simsplit | -972576 | 1945193 |
| ancmig_adj_1 | -984705 | 1969431 |
| refugia_adj_5_full_2 | -996910 | 1993887 |
| ancmig_adj_2 | -1106291 | 2212596 |
| sim_split_no_mig_size | -1479848 | 2959714 |
| split_nomig | -1532011 | 3064034 |
| sim_split_no_mig | -1542949 | 3085908 |

**Table S2: Demographic model selection.** Log-likelihoods and Akaike Information Criterion (AIC) values for the 18 demographic models tested, following model optimisation.
